# Understanding the role of memory re-activation and cross-reactivity in the defense against SARS-CoV-2

**DOI:** 10.1101/2021.07.23.453352

**Authors:** Viola Denninger, Catherine K. Xu, Georg Meisl, Alexey S. Morgunov, Sebastian Fiedler, Alison Ilsley, Marc Emmenegger, Anisa Y. Malik, Monika A. Piziorska, Matthias M. Schneider, Sean R. A. Devenish, Vasilis Kosmoliaptsis, Adriano Aguzzi, Heike Fiegler, Tuomas P. J. Knowles

## Abstract

Recent efforts in understanding the course and severity of SARS-CoV-2 infections have highlighted both potential beneficial as well as detrimental effects of cross-reactive antibodies derived from memory immunity. Specifically, due to a significant degree of sequence similarity between SARS-CoV-2 and other members of the coronavirus family, memory B-cells that emerged from previous infections with endemic human coronaviruses (HCoVs) could be re-activated upon encountering the newly emerged SARS-CoV-2, thus prompting the production of cross-reactive antibodies. Understanding the affinity and concentration of these potentially cross-reactive antibodies to the new SARS-CoV-2 antigens is therefore particularly important when assessing both existing immunity against common HCoVs and adverse effects like antibody-dependent enhancement (ADE) in COVID-19. However, these two fundamental parameters cannot easily be deconvoluted by surface-based assays like enzyme-linked immunosorbent assays (ELISAs) which are routinely used to assess cross-reactivity.

Here, we have used microfluidic antibody-affinity profiling (MAAP) to quantitatively evaluate the humoral immune response in COVID-19 convalescent patients by determining both antibody affinity and concentration against spike antigens of SARS-CoV-2 directly in nine convalescent COVID-19 patient and three pre-pandemic sera that were seropositive for common HCoVs. All 12 sera contained low concentrations of high affinity antibodies against spike antigens of HCoV-NL63 and HCoV-HKU1, indicative of past exposure to these pathogens, while the affinity against the SARS-CoV-2 spike protein was lower. These results suggest that cross-reactivity as a consequence of memory re-activation upon an acute SARS-CoV-2 infection may not be a significant factor in generating immunity against SARS-CoV-2.

## Introduction

Despite the increasing availability of diagnostic tests, antibody treatments and vaccines, the COVID-19 pandemic still presents a major challenge to governments and healthcare systems all over the world, requiring a functional assessment of the immune response to SARS-CoV-2. Research focuses increasingly around the role of acute increased production of pro-inflammatory cytokines and the recall of memory T- and B-cells which were formed during previous infections with other coronaviruses^1–4^. In contrast to SARS-CoV-2, common human coronaviruses (HCoVs), although omnipresent and recurrent, generally cause mild disease only^5^. HCoVs can be detected throughout all age-groups with most individuals seroconverting during childhood^6, 7^. Re-infection subsequently leads to antibody affinity maturation, resulting in a natural build-up of protective memory T- and B-cells as well as long lived plasma cells (LLPCs) that in some cases can persist up to a year^8, 9^. In the event of the emergence of new but highly homologous coronavirus species, memory B-cells might recognize the antigens of the new species, re-activate and accelerate the immune response to the new virus, prompting the production of cross-reactive antibodies^10^.

Although immunity derived from previous infections with similar viruses is generally advantageous, cross-reactive antibodies can cause antibody-dependent enhancement (ADE) of the viral infection in a subset of individuals^10, 11^. In these cases, cross-reactive antibodies, although able to recognize and bind the new viral antigens, not only fail to neutralize the virus due to sub-neutralizing concentrations, insufficiently high affinity, or binding to an inappropriate region, but more importantly facilitate its reproduction via an Fc-receptor mediated uptake into the host cell. Cross-reactivity mediated ADE has been shown to elevate the disease severity of infections with flaviviruses, especially for Dengue and Zika viruses, and is therefore a key consideration in the development of vaccines and therapies for COVID-19^12–14^.

Cross-reactivity of antibodies is traditionally assessed using enzyme-linked immunosorbent assays (ELISAs). These assays report titers only and cannot readily resolve the fundamental parameters of antibody responses, namely affinity and concentration^15^, which are crucial for monitoring the effectiveness of humoral immunity after natural infection and after vaccination.

Here, we have used microfluidic antibody-affinity profiling (MAAP)^15–17^ to quantify the cross-reactivity of antibodies against SARS-CoV-2 S1, S2 and the receptor binding domain (RBD) in nine convalescent COVID-19 patient samples and three pre-pandemic controls, and compared our results to standard ELISA assays. While MAAP matched the ELISA data qualitatively, crucially it enabled the simultaneous and independent determination of antibody affinities and concentrations to reveal interindividual variation of antibody responses to SARS-CoV-2 not detectable by ELISA.

Interestingly, testing all 12 serum samples in a reciprocal approach for antibodies against the corresponding antigens of two common HCoVs, namely HCoV-NL63 and HCoV-HKU1, revealed high antibody affinity and low concentrations across all samples tested. This indicates the presence of matured antibodies against both common HCoVs, rather than cross-reactive antibodies that were derived from the immune response against SARS-CoV-2. This observation was further confirmed by single point competition assays using the SARS-CoV-2 RBD and HCoV-NL63 RBD antigens.

Our data therefore suggests no significant cross-reactivity of antibodies against common HCoVs to SARS-CoV-2 and vice versa, indicating that B-cell mediated immunological memory from past infections with common HCoVs is unlikely to have a significant role in SARS-CoV-2 specific immune response.

## Results and Discussion

### Quantification of the immune response to SARS-CoV-2 in convalescent COVID-19 patient sera

Before assessing cross-reactivity, we first quantified the immune response to SARS-CoV-2 in nine seropositive convalescent COVID-19 patient samples using MAAP^16^. In addition to profiling the receptor binding domain (RBD), we extended the assay to also include the spike S1 and S2 subunits as the spike S2 subunit had previously been reported to comprise regions of high homology with other human coronaviruses and therefore may confer a higher probability of cross-reactivity to antibodies from previous infections^18^.

MAAP against the SARS-CoV-2 RBD region revealed *K_D_* values ranging from 1–28 nM which is consistent with previously reported values in patients with acute COVID-19 infections^16^ (**Figure 1A**). Interestingly, serum samples **4** and **5** not only contained high concentrations of antibody binding sites against the SARS-CoV-2 RBD region (serum **4**: 169 nM (CI 95%: 123–208) and serum **5**: 231 nM (CI 95%: 152–285)), but also against SARS-CoV-2 spike S1 (serum **4**: 100 nM (CI 95%: 81–123), and serum **5**: 480 nM (CI 95%: 351–533)) and SARS-CoV-2 spike S2 (serum **4**: 90 nM (CI 95%: 66–100), and serum **5**: 351 nM (CI 95%: 231–390)). This may be due to earlier sampling in these patients after recovery from COVID-19 compared to the other seven sera tested, so that the antibody concentration had not declined significantly yet.

**Figure 1.**
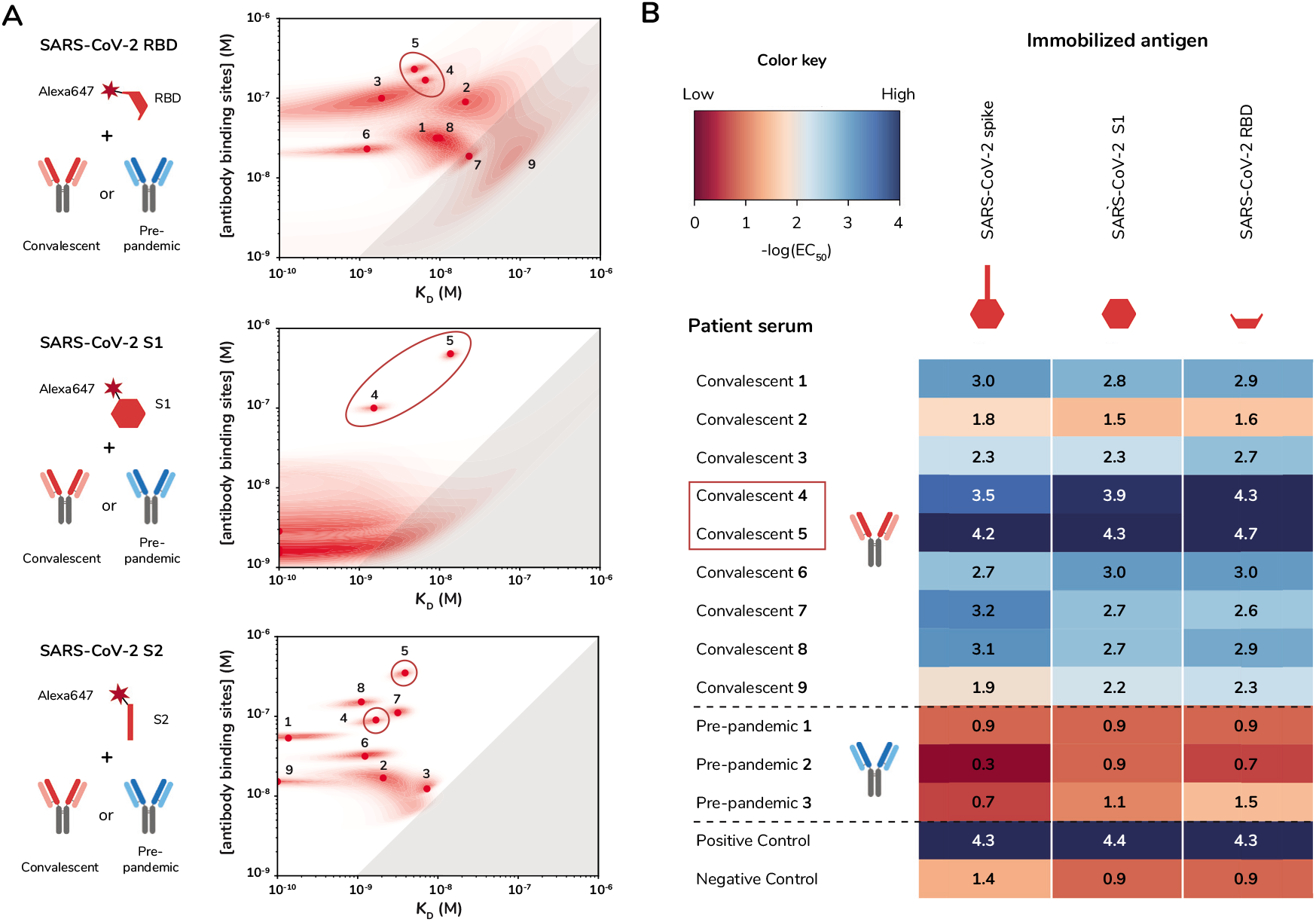
Microfluidic antibody-affinity profiling (MAAP) of nine convalescent COVID-19 patient and three pre-pandemic sera. **(A)** Probability density plots of MAAP against fluorescently labeled SARS-CoV-2 RBD, spike S1 and spike S2. The graphs show the affinity (*K_D_*) and the molar concentration of antibody binding sites for each convalescent serum sample. No binding could be detected in the three pre-pandemic sera. Points correspond to the maximum a posteriori values in the two-dimensional posterior probability distribution, and shaded regions to the probability density. Grey shaded regions indicate the area of non-binding for samples with [antibody binding sites] <2*K_D_*. **(B)** ELISA −log (EC_50_) values for the same nine convalescent COVID-19 patient samples and three pre-pandemic sera (**1**: NL63+, 229E+; **2**: NL63+, 229E+, OV43+, HKU1+; **3**: 229E+). A SARS-CoV-2 seronegative sample and a SARS-CoV-2 seropositive sample served as negative and positive controls, respectively. Immobilized antigens are SARS-CoV-2 spike ectodomain, spike S1 and the RBD. Detection was performed with fluorescently labeled anti-human IgG antibodies. **(A)** and **(B)** Red boxes/ circles indicate convalescent sera **4** and **5** which have the strongest immune response to all SARS-CoV-2 spike subunits.

All nine convalescent serum samples contained antibodies with high affinity to the S2 subunit of SARS-CoV-2 spike with the majority of the *K_D_* values in the lower nanomolar to even sub-nanomolar range. There was no correlation between the concentration of antibody binding sites against the S2 subunit and the RBD region, supporting previous reports that indicate highly variable immune responses between individual patients towards these two different antigens^18, 19^.

While antibodies in all nine convalescent sera were found to bind tightly to the S2 subunit, we found considerable variation in affinity and concentration of serum antibodies when using MAAP against the S1 subunit. Two out of nine convalescent serum samples exhibited tight binding (*K_D_* <10 nM) but low concentrations of antibody binding sites against the spike S1 subunit (sera **6** and **7**). Moreover, for serum **1** no affinity could be determined potentially due to low antibody concentration or affinity. A further four samples did not show any binding to spike S1 (sera **2, 3, 8** and **9**), potentially due to a closed conformational state of the S1 subunit, obscuring crucial epitopes from antibody binding as previously observed for HCoV-229E^20–22^ (**Suppl. Table 1**).

To validate these results by use of an independent assay, we also subjected the convalescent serum samples to a standard ELISA. As ELISAs employ antigens that are immobilized to a surface rather than measuring the interaction in solution, the assay output is strongly dependent on the combination of antibody affinity, avidity and concentration. However, the assay does not report on these crucial parameters individually. While the ELISA qualitatively corroborated the MAAP results and identified sera **4** and **5** as the strongest responders to all three SARS-CoV-2 antigens (**Figure 1B**), our assay in addition deconvoluted the respective contributions of antibody affinity and concentration towards the qualitative EC_50_ values determined by the ELISA assay.

### Highly specific antibodies against HCoV-NL63 and HCoV-HKU1 spike S1 and RBD are present at low concentrations in convalescent COVID-19 and pre-pandemic sera

Receptor-mediated cell entry of coronaviruses typically depends on the interaction of the spike protein with a target receptor that is specific to each coronavirus. Opposed to that, it was shown that neutralizing antibodies targeting the spike proteins of SARS-CoV and MERS-CoV can instead facilitate Fc-receptor mediated cell entry and virus propagation^11, 23^, a mechanism typical for ADE. This raises the possibility of cross-reactive anti-spike antibodies influencing acute SARS-CoV-2 infections or vaccinations. Indeed, several recent studies have investigated cross-reactivity between SARS-CoV-2 and HCoV antibodies, with particular focus on antibody binding to the spike protein and RBD^4, 19, 24, 25^. The studies found that most pre-pandemic sera contained antibodies that bind to the spike protein of all common HCoVs, while binding to the spike protein of SARS-CoV-2 was observed less frequently. Conversely, convalescent COVID-19 patient sera seemed to show cross-reactivity to the spike protein of all common HCoVs, albeit to different degrees^4, 16, 19^.

To test these observations in our system, we used MAAP against the RBD and S1 subunit of HCoV-NL63 and HCoV-HKU1. Even though sequence alignment of the full length spike proteins of both HCoVs with the SARS-CoV-2 spike protein revealed ~30% sequence identity at the amino acid level, the betacoronavirus HCoV-HKU1 was previously reported to be more readily recognized by cross-reactive antibodies in sera of recovered COVID-19 patients^4, 19^. As HCoV-HKU1 binds to a different cell receptor however, structural differences are likely in the RBD region, the most crucial antigen for binding of potentially neutralizing, cross-reactive antibodies. HCoV-NL63 on the other hand, like SARS-CoV-2, binds to the cell-surface receptor ACE2 to enter the host cell, resulting in similarities in the RBD^26^.

As an initial proof of concept, we determined the binding affinity of a recombinantly expressed neutralizing anti-SARS-CoV-2 RBD antibody to the S1-subunits of HCoV-NL63 and HCoV-HKU1, respectively. To do so, we measured the hydrodynamic radius, *R_h_*, of the fluorescently labeled S1-subunits (**Figures 2A and B**) at increasing concentrations of the anti-SARS-CoV-2 RBD antibody. Whereas the antibody readily bound SARS-CoV-2 S1 with a *K_D_* of ~5 nM (data not shown), the size of both labeled HCoV S1 subunits remained unchanged, even up to antibody concentrations of 500 nM, indicating the absence of binding. As a control, the same assay was performed titrating an anti-HKU1 antibody against fluorescently labeled HCoV-HKU1 spike S1 (**Figure 2C**), clearly showing a high affinity of the antibody to the labeled HCoV-HKU1-antigen (*K_D_* <100 pM). These results indicate that there is no substantial cross-reactivity with the S1 subunits of HCoV-HKU1 and HCoV-NL63 for the anti-SARS-CoV-2 RBD antibody investigated here, even though HCoV-NL63 and SARS-CoV-2 share ACE2 as their target receptor^27^.

**Figure 2.**
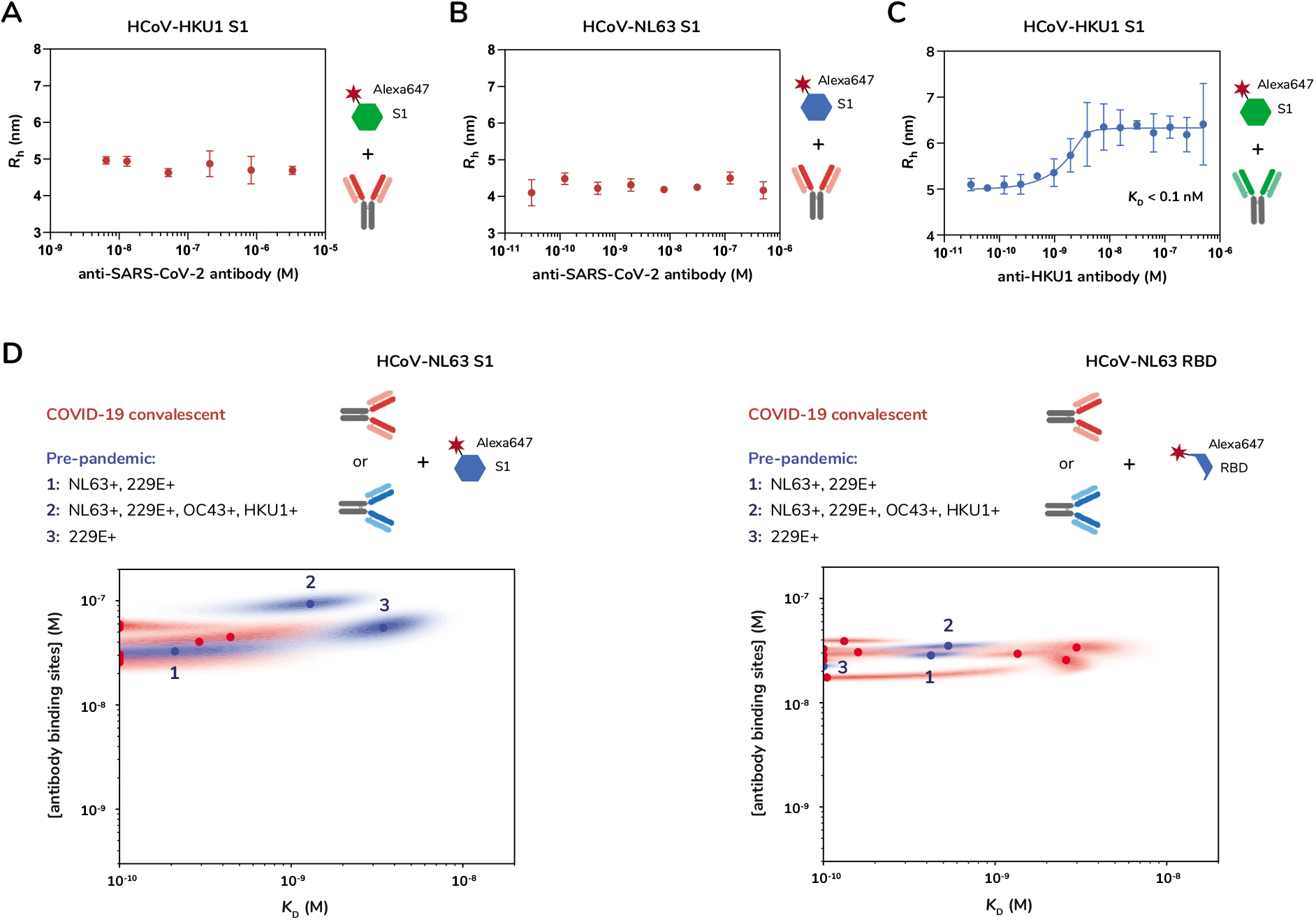
Microfluidic antibody-affinity profiling against HCoV-NL63 spike S1 and RBD to establish cross-reactivity of antibodies in convalescent COVID-19 serum. **(A)** and **(B)** Equilibrium binding curves of a neutralizing SARS-CoV-2 antibody against 10 nM fluorescently labeled spike S1 from **(A)** HCoV-HKU1 or **(B)** HCoV-NL63, respectively. The hydrodynamic radii (*R*_h_) of the free labeled spike proteins did not increase upon addition of the antibody, indicating the absence of binding. **(C)** Equilibrium binding curve of an anti-HKU1 antibody against 10 nM fluorescently labeled spike S1 of HCoV-HKU1 showed very tight binding with a *K_D_* below 0.1 nM. The *K_D_* was determined by non-linear least squares fitting using Equation 1. **(D)** Probability density plots of MAAP against fluorescently labeled HCoV-NL63 spike S1 and HCoV-NL63 RBD. The graphs show the affinity (*K_D_*) and the molar concentration of antibody binding sites for each of the convalescent COVID-19 (red) and the pre-pandemic sera (blue) whereby pre-pandemic sera **1** and **2** were found to be seropositive for HCoV-NL63. Points correspond to the maximum a posteriori values in the two-dimensional posterior probability distribution, and shaded regions to the probability density.

After this initial proof of concept, we tested the previously mentioned nine convalescent COVID-19 patient sera (**Figure 2D**) together with three pre-pandemic sera, one of which was previously confirmed to be seropositive for HCoV-NL63 and HCoV-HKU1 (pre-pandemic serum **2**), and another one to be seropositive for HCoV-NL63 only (pre-pandemic serum **1**).

In all nine convalescent sera we detected moderate concentrations of antibody binding sites for HCoV-NL63 RBD and S1, ranging from 17 to 40 nM and from 27 to 60 nM (**Figure 2C and Suppl. Table 2**). Concentrations of antibody binding sites against the respective HCoV-HKU1 antigens were even lower with 3.4–22 nM for HCoV-HKU1 RBD and 3.5–37 nM for HCoV-HKU1 S1 (**Suppl. Figure 1 and Suppl. Table 3**). With concentrations >90 nM, only one pre-pandemic serum (pre-pandemic serum **2**) exhibited higher levels of antibody binding sites against the S1 subunits of HCoV-NL63 and HCoV-HKU1, indicative of an increased antibody production likely in response to a recent infection with both viruses.

Intriguingly, with *K_D_* values <3.6 nM, all sera contained antibodies with very high affinities against the two HCoV-NL63 antigens. Furthermore, affinities against HCoV-HKU1 RBD were in the sub-nanomolar range throughout all samples that were subjected to the MAAP assay. The high specificity towards HCoV-NL63 and HCoV-HKU1 epitopes indicates that these antibodies were not produced in response to SARS-CoV-2, but were rather derived from memory immunity. Based on our observations as well as previously published reports^4, 6, 28^, we hypothesize that the antibodies we detected with use of our MAAP assay are part of a general background immunity of high affinity circulating serum antibodies against HCoV-NL63 and HCoV-HKU1 that were produced in response to previous infections.

### Pre-pandemic sera show no significant cross-reactivity against SARS-CoV-2 spike S1, S2 and RBD

Given the ubiquitous prevalence and high re-infection rate of common HCoVs^29^, it is likely that most individuals will have preexisting antibodies deriving from immunological memory circulating in the bloodstream at concentrations comparatively lower than those in response to an acute infection^9^. If cross-reactive to SARS-CoV-2 antigens, these preexisting antibodies could not only impact the accuracy of diagnostic tests for SARS-CoV-2 infections, but could also present a risk for ADE, and thus potentially affect the severity of the immune response after infection. It is therefore crucial to further examine the ability of anti-HCoV antibodies to recognize and bind SARS-CoV-2 antigens. To probe for the cross-reactive potential of these antibodies against the corresponding SARS-CoV-2 epitopes, we performed MAAP assays against the three SARS-CoV-2 antigens, spike S1, spike S2 and RBD in the three pre-pandemic sera.

Interestingly, affinity profiling did not reveal any detectable binding of preexisting antibodies to SARS-CoV-2 RBD, spike S1 or spike S2 in the three pre-pandemic sera (**Figure 1A and Suppl. Table 1**).

To further investigate, we then assessed cross-reactivity directly via a competition assay based on the principle described in Schneider *et al*^16^ and Fiedler *et al^17^* (**Figure 3A**). For this approach, a quantity of each serum corresponding to 20 nM antibody binding sites was mixed with 10 nM labeled HCoV-NL63 RBD, to allow at least 80% of the labeled antigen to be bound by antibodies. After equilibration, the mixture was subjected to microfluidic diffusional sizing, and complex formation was confirmed by the increase of the apparent *R*_h_. For competition, an excess of 250 nM unlabeled SARS-CoV-2 RBD was added to the HCoV-NL63 RBD–antibody complex, and *R*_h_ was measured after establishment of binding equilibrium. If antibodies bound to the RBD of HCoV-NL63 were cross-reactive, the excess unlabeled SARS-CoV-2 RBD would outcompete the binding, resulting in an unbound HCoV-NL63 RBD and therefore a decrease in *R*_h_ compared to that of the antibody complexed HCoV-NL63 RBD.

**Figure 3.**
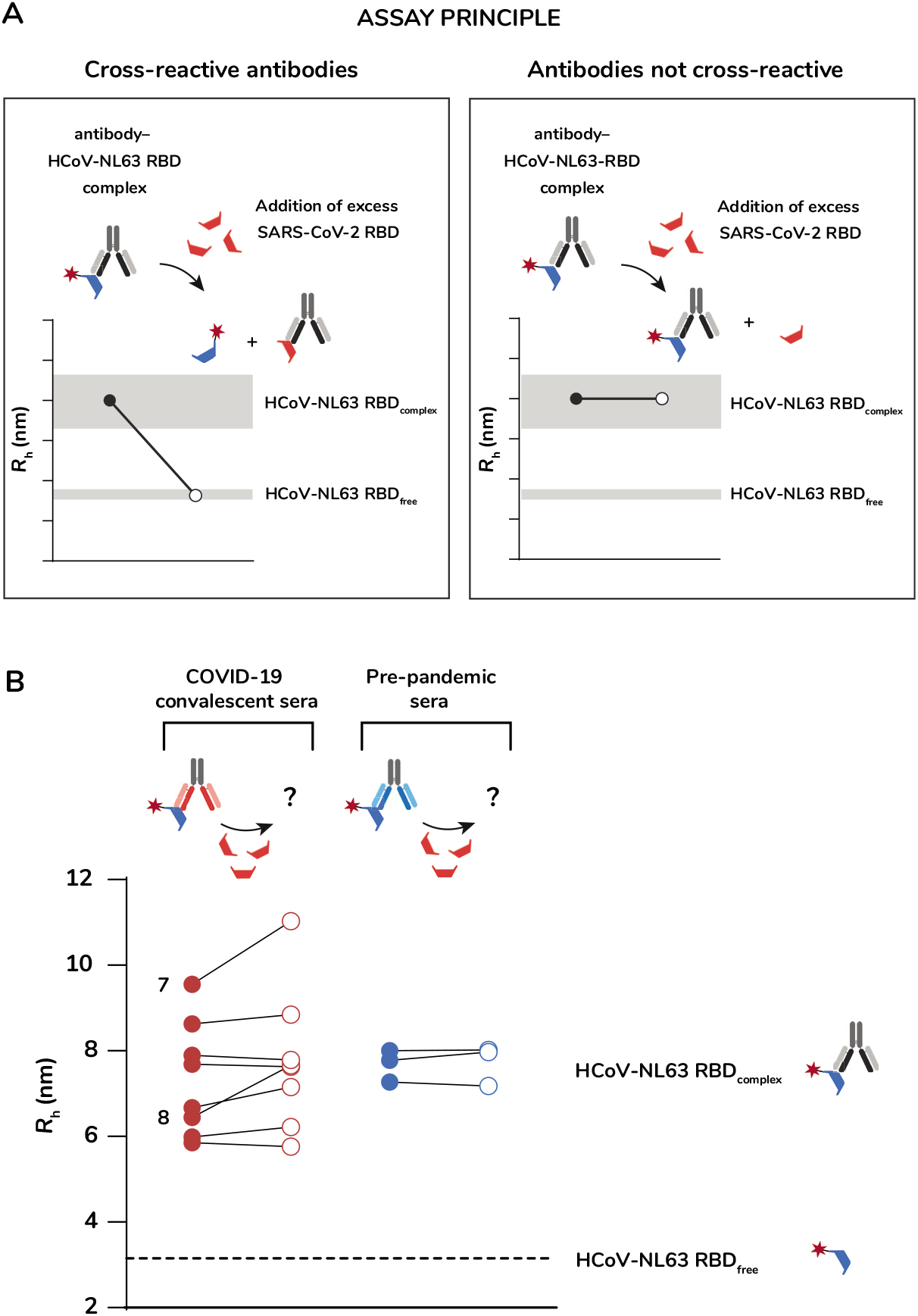
**(A)** Schematic of SARS-CoV-2 - HCoV-NL63 cross-reactivity competition assay. 10 nM fluorescently labeled HCoV-NL63 RBD was mixed either with buffer or with patient serum, incubated 1 h and subjected to microfluidic diffusional sizing, to determine the size of the free labeled RBD (*R_h,free_*) and the size of the immune-complex (*R_h,complex_*). For binding competition, 250 nM unlabeled SARS-CoV-2 RBD was added to the mixture of 10 nM HCoV-NL63 RBD and patient serum and incubated for 1 h before measuring the hydrodynamic radius (*R*_h_). If the serum contains cross-reactive antibodies, a size decrease is observed, as the antibodies will bind to the excess of unlabeled RBD (left box). If the antibodies in the serum are not cross-reactive, they remain bound to the labeled HCoV-NL63 RBD and the *R_h_* of the immuno-complex will stay constant (right box). **(B)** SARS-CoV-2-HCoV-NL63 cross-reactivity competition assay in convalescent COVID-19 sera and pre-pandemic sera. Filled circles (o) indicate the size of the HCoV-NL63 RBD–anti-body immune-complex. Empty circles (o) indicate the size of HCoV-NL63 RBD after addition of excess unlabeled SARS-CoV-2 RBD. (Red) Convalescent COVID-19 sera, (Blue) pre-pandemic sera. No decrease of *R_h_* upon addition of the unlabeled SARS-CoV-2 RBD could be observed in any of the sera, indicating no cross-reactivity of an-ti-NL63 antibodies with SARS-CoV-2 RBD.

Using this approach, we again tested the nine convalescent COVID-19 as well as the three HCoV pre-pandemic sera. We found that even upon addition of excess unlabeled SARS-CoV-2 RBD, the *R*_h_ did not decrease in any of the samples, indicating that all antibodies stayed tightly bound in a complex with labeled HCoV-NL63 RBD, and thus were not cross-reactive with SARS-CoV-2 RBD (**Figure 3B**). Two convalescent COVID-19 serum samples however, serum **7** and serum **8**, presented an interesting exception, as their complex size increased upon addition of unlabeled SARS-CoV-2 RBD. This was likely due to unsaturated antibody binding sites that were still free to bind some of the SARS-CoV-2 RBD protein, but with a low affinity unable to outcompete the interaction with labeled HCoV-NL63 RBD (**Suppl. Tables 1 and 2**). Additionally, we observed greater variations of initial HCoV-NL63 RBD–antibody complex sizes within the pool of convalescent COVID-19 serum samples (*R*_h_= 5.9–9.6 nm) compared to the pre-pandemic sera (*R*_h_= 7.3–8.0 nm). This observation is in line with the fact that our cohort derived from recently COVID-19 recovered patients which contained antibody types of different sizes, namely IgG and IgM.

Based on these results, even though we cannot fully exclude cross-reactivity against the SARS-CoV-2 RBD, we would expect cross-reactive antibodies to exhibit at least 10-fold lower affinity to the SARS-CoV-2 RBD region compared to its original antigen.

## Conclusions

Protective and adverse effects of antibody cross-reactivity can influence the humoral immune response of an individual to an acute infection, vaccination, or treatment. In this study, we have employed microfluidic antibody-affinity profiling (MAAP) to quantitatively investigate cross-reactivity against SARS-CoV-2 and two common coronaviruses, HCoV-NL63 and HCoV-HKU1, in nine convalescent COVID-19 and three pre-pandemic sera. This enabled us to quantify the affinity and concentration of antibodies against different spike subunits of the three coronaviruses, allowing a differentiation between antibodies raised in response to an acute infection and those originating from immunological memory. Affinity profiling against different SARS-CoV-2 spike antigens revealed a wide range of antibody affinities and concentrations in all convalescent sera, especially against the RBD and the S1 subunit of SARS-CoV-2. In our cohort, this variability suggests a *de novo* synthesis of antibodies in response to a SARS-CoV-2 infection, further supported by the fact, that we did not find evidence for significant cross-reactivity of antibodies against SARS-CoV-2 spike antigens in pre-pandemic, HCoV-seropositive sera.

This contrasts with the antibody affinities and concentrations that were found when profiling the same 12 sera against spike S1 and RBD from HCoV-NL63 and HCoV-HKU1. With the exception of one pre-pandemic serum that was seropositive for all common HCoVs and thus showed elevated levels of antibodies against HCoV-NL63 and HCoV-HKU1, all sera tested contained antibodies with high affinity, but at relatively low antibody concentrations. Consistent with epidemiological studies on infection patterns of HCoV-NL63 and HCoV-HKU1^30^, this suggests the existence of a low amount of circulating antibodies against the RBD and the S1 subunit of the two HCoVs, that already underwent affinity maturation in response to re-occurring infections, rather than being the result of cross-reactivity induced re-call of memory B-cells^28^.

By deconvoluting the humoral immune response into affinity and concentration of serum antibodies, MAAP generates an individual fingerprint for each sample, providing crucial insights into the immunological landscape of *de novo* and reoccurring infections. This unique quantitative approach therefore delivers new prospects to monitor immune induction following a vaccination or a treatment more precisely, helping to evaluate the efficacy of antibody production and to assess the potential risks of humoral memory.

## Materials and Methods

### Sequence alignment

Sequence alignment of coronavirus full length spikes was performed using the Clustal Omega Sequence Alignment Tool. Input sequences were HCoV-NL63 (UniProt ID: Q6Q1S2), HCoV-HKU1 (UniProt ID: Q5MQD0) and SARS-CoV-2 (UniProt ID: P0DTC2).

### Origin of serum samples

All serum samples from COVID-19 convalescent patients as well as the pre-COVID sera were purchased from BioIVT. Pre-COVID sera were seropositive for the following coronaviruses: serum **1** (HCoV-229E, HCoV-NL63), serum **2** (HCoV-229E, HCoV-NL63, HCoV-OC43, HCoV-HKU1) and serum **3** (HCoV-229E). BioIVT sought informed consent from each subject, or the subjects legally authorized representative and appropriately documented this in writing. All samples are collected under IRB-approved protocols.

### ELISA assay

ELISA-based serology was carried out as previously described^24, 31^, with minor modifications. High-binding 384-well plates (Perkin Elmer, SpectraPlate 384 HB) were coated with 20 μL 1 μg/mL spike ectodomain (Lausanne, EPFL SV PTECH PTPSP), RBD (Lausanne, EPFL SV PTECH PTPSP), or spike S1 domain (S1N-C52H4, Acro Biosystems) in PBS at 37 °C for 1 h, followed by 3 washes with PBS 0.1% Tween-20 (PBS-T) using a Biotek EL406 plate washer and blocked with 40 μL 5% milk in PBS-T for 1.5 h. Serum samples were serially diluted (range: 0.02 – 1.2×10^−6^) in sample buffer (1% milk in PBS-T) and added to wells (20 μL/well). After sample incubation for 2 h at RT, the wells were washed five times with wash buffer and the presence of IgGs directed against above-defined SARS-CoV-2 antigens was detected using an HRP-linked anti-human IgG antibody (Peroxidase AffiniPure Goat Anti-Human IgG, Fcγ Fragment Specific, Jackson, 109-035-098, at 1:4000 dilution in sample buffer) at 20 μL/well. The incubation of the secondary antibody for one hour at RT was followed by three washes with PBS-T, the addition of TMB, an incubation of five minutes at RT, and theaddition of 0.5 M H_2_SO_4_. The plates were centrifuged after all dispensing steps, except for the addition of TMB. The absorbance at 450 nm was measured in a plate reader (Perkin Elmer, EnVision) and the inflection points of the sigmoidal binding curves (p(EC_50_) or −log(EC_50_) values of the respective sample dilution) were determined using the custom designed fitting algorithm previously reported^24^. The p(EC_50_) values for all samples and antigens were visualized using the ggplot2 package in R.

### Fluorescent labelling of proteins

SARS-CoV-2 S1 protein (S1N-C52H4), SARS-CoV-2 S protein RBD (SPD-C52H3) and SARS-CoV-2 S2 protein (S2N-C52H5) were purchased from Acro Biosystems. HCoV-NL63 S1 protein (40600-V08H) and HCoV-HKU1 S1 protein (40021-V08H) were purchased from Sino Biological. HCoV-NL63 Spike RBD (10605-CV) and HCoV-HKU1 Spike RBD (10600-CV) were obtained from R&D Systems.

All proteins were reconstituted according to manufacturers’ instructions. For fluorescent labeling, the protein was diluted into labeling buffer (1 M NaHCO3 pH 8.3) to give a final concentration of 0.2 M NaHCO_3_, followed by the addition of Alexa Fluor™ 647 NHS ester (A2006, Thermo Scientific) at a molar dye-to-protein ratio of 3:1. Following incubation overnight at 4 °C, the labeled proteins were purified by size exclusion chromatography (SEC) on an ÄKTA pure system (Cytiva) with phosphate buffered saline (PBS) at pH 7.4 (P4417, Merck) as elution buffer. For SEC-purification of RBD proteins a Superdex 75 Increase 10/300 GL column was used, the larger S1 and S2 proteins were purified using a Superdex 200 Increase 3.2/300 column (Cytiva). Pooled fractions that correspond to the labeled protein were concentrated, snap frozen and stored at −80 °C with 10% v/v glycerol as a cryoprotectant.

### Equilibrium affinity binding curves of recombinant antibodies against S1 proteins of HCoV-HKU1 and HCoV-NL63

For affinity measurements of anti-human coronavirus spike glycoprotein HKU1 (40021-MM07-100, Sino Biological) to HCoV-HKU1 S1 protein, the antibody was reconstituted according to the manufacturer’s instructions and diluted into PBS, containing 0.05% Tween 20 and 5% human serum albumin (HSA), to achieve a two-fold concentration series ranging from 60 pM to 1 μM. Antibody dilutions were subsequently mixed in a 1:1 ratio with a 20 nM solution of Alexa Fluor 647 labeled HCoV-HKU1 S1, to obtain a final concentration of 10 nM.

For cross-reactivity measurements, anti-SARS-CoV-2 neutralizing antibody (SAD-S35, Acro Biosystems) was reconstituted according to the manufacturer’s instructions. A four-fold dilution series, ranging from 60 pM to 1 μM was prepared by diluting into PBS, containing 0.05% Tween 20 and 5% human serum albumin (HSA). Antibody dilutions were subsequently mixed in a 1:1 ratio with a 20 nM solution of Alexa Fluor 647 labeled HCoV-HKU1 S1 or Alexa Fluor 647 labeled HCoV-NL63 S1 respectively, to obtain a final concentration of 10 nM.

All samples were incubated for 1 h at 4 °C prior to measurement and kept at 4 °C throughout the experiment. Measurements were performed in triplicate using the 1.5–8 nm size-range setting on the Fluidity One-W Serum (Fluidic Analytics). The binding affinity, *K_D_*, was generated by non-linear least squares fitting to Equation 1, leaving all parameters to be fit, but constrained to be >0.

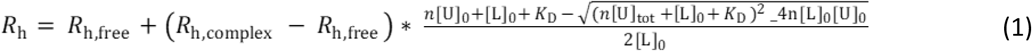

with *R*_h_, *R*_h,free_, and *R*_h,complex_ being the average hydrodynamic radii of the reaction at equilibrium, of the unbound labeled S1 protein and of the complex of labeled S1 with unlabeled antibody, respectively. The parameters [U]_0_ and [L]_0_ are total concentrations of unlabeled antibody and labeled S1, respectively, and n is the number of binding sites per unlabeled antibody.

### Microfluidic antibody-affinity profiling of serum antibodies (MAAP)

Measurements of serum samples were performed as reported previously^16^. For the MAAP measurements, varying fractions of COVID-19 convalescent or pre-COVID serum samples were combined with Alexa Fluor 647 labeled antigen of varying concentrations ranging between 1 nM and 250 nM. Buffer was added to give a constant volume of 20 μL for each sample, followed by an incubation for 1 h at 4 °C. For MAAP against SARS-CoV-2 RBD, HCoV-NL63 RBD or HCoV-HKU1 RBD, PBS containing 0.05 % Tween 20 (pH 7.4) was used as a buffer. For profiling of serum against SARS-CoV-2 S1, SARS-CoV-2 S2, HCoV-NL63 S1 and HCoV-HKU1 S1, the buffer additionally contained 5% human serum albumin (HSA). Because of limited sample availability only a subset of the nine convalescent serum samples could be affinity profiled against HCoV-HKU1 RBD and HCoV-HKU1 S1 (**Suppl. Figure 1 and Suppl. Table 3**).

Following incubation, the hydrodynamic radius, *R*_h_, of the serum antibody – antigen complex was determined by microfluidic diffusional sizing using a Fluidity One-W Serum. Additionally, each serum was measured separately at a concentration of 100%, 50%and 25%, to allow correction of titration data-points for the autofluorescence of the individual serum.

The binding affinities and concentrations of antigen-specific antibodies in serum samples were determined by monitoring the fraction of labeled antigen that diffused into the distal chamber of the microfluidic device. Following background correction of samples to account for autofluorescence signal arising from the serum, this fraction can be determined by:

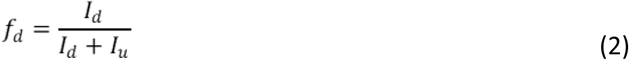

Given that we monitor a mixture of free and bound (B) labeled species (L), *I*_d_ and *I*_u_ can be expressed as:

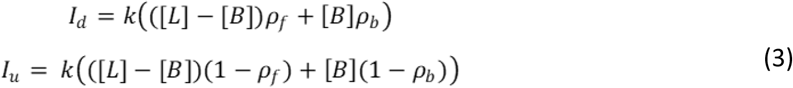

where *k* is a constant that relates the concentration of L to the observed fluorescence intensity, [B] is the equilibrium concentration of bound L, and *ρ_f_* and *ρ_b_* are the fractions of free and bound labeled antigen that diffuse across to the distal channel, respectively. *f_d_* can thus be expressed as follows:

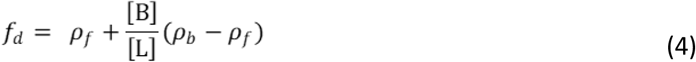

By solving the binding equilibrium equation, [B] is given by:

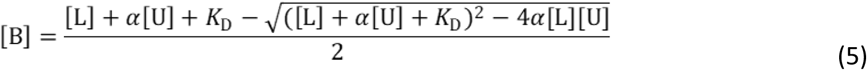

where [U] is the total concentration of antibody binding sites in the serum, and α is the fractional concentration of serum in the binding measurement.

Bayesian inference was used to determine *K_D_* and [U], with a prior that was flat in logarithmic space for *K_D_* and [U], and flat in linear space for *ρ*_f_ and *ρ*_b_. The likelihood was a Gaussian with mean *f_d_* and its standard deviation defined as the standard deviation of all measurements of each labeled antigen.

As serum samples were derived from convalescent patients, potentially containing different classes of antibodies with variable numbers of binding sites (IgG = 2, IgM = 10), antibody concentration is expressed in terms of binding sites, rather than molecules.

### SARS-CoV-2 RBD and HCoV-NL63-RBD competition assay

10 nM Alexa Fluor 647 labeled HCoV-NL63 RBD in PBS containing 0.05% Tween 20 was combined with serum sample corresponding to a final anti-NL63 RBD antibody concentration of 20 nM (based on previous MAAP of each serum sample). For competition, unlabeled SARS-CoV-2 RBD was added at a concentration of 250 nM. Samples were incubated 1 h at 4 °C prior to equilibrium binding measurements using the Fluidity One-W Serum. Labeled HCoV-NL63 RBD in buffer, and a mixture of labeled HCoV-NL63 RBD with serum only served as controls. All measurements were performed in triplicate. For background correction, serum was diluted in PBS containing 0.05% Tween 20, at the same v/v concentration as used in the assay and measured separately. Background subtraction was applied to individual datapoints obtained in the assay.

### Quality control

As part of constructing the dataset used in this research, we developed a series of procedures that we jointly refer to as “quality control (QC) pipeline”. Firstly, they serve as simple sanity checks to ensure that data processing and model fitting was performed correctly. Secondly, they help identify problematic measurements, samples, or model fits. Based on this, decisions to re-collect or exclude parts of the data can be made. Finally, if assumptions about relationships in the dataset are appropriately tested, information can be shared between experiments and samples in a robust way, which improves model fitting. The detailed discussion of these methods is outside the scope of this publication; however, we are providing it in the form of a supplementary to this work, to serve as a guide for anyone performing similar analyses (**Suppl. Material 1**). When applying the described procedures to the present dataset, we identified a small number of outlier measurements. These were removed during the model fitting and data analysis step. Otherwise, the dataset passed all QC tests.

## Funding Acknowledgements

Work performed by AA and ME was funded by institutional core funding by the University of Zurich and the University Hospital of Zurich, Driver Grant 2017DRI17 of the Swiss Personalized Health Network (SPHN), Distinguished Scientist Award of the NOMIS Foundation, a Grant of the European Research Council (ERC Prion2020 670958), and an Innovation Fund of the University Hospital Zurich. V.K. was funded by an NIHR fellowship (PD-2016-09-065) and acknowledges support as a PI Terasaki Scholar. T.P.J.K. is grateful for financial support by the Biotechnology and Biological Sciences Research Council and the European Research Council.

## Author Contributions

Microfluidic study design and measurements: VD, SF, AI, AYM, MAP, MMS, SRAD, HF. Data Analysis: VD, CKX, GM, ASM, SF. Data Quality Control: ASM. ELISA design, measurements, and data analysis: ME, AA. Study advice: AA, VK. Study supervision: HF, TPJK. Manuscript writing: VD, MMS, HF.

