## Supplementary Tables and Figures for "Understanding the role of memory re-activation and cross-reactivity in the defense against SARS-CoV-2"

### Supplementary Figure 1 and Tables 1 to 3

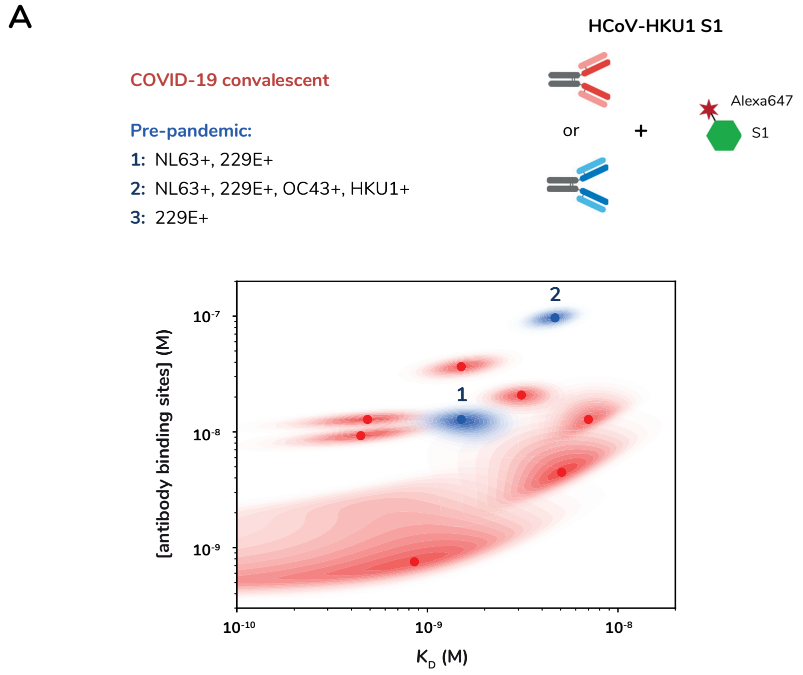


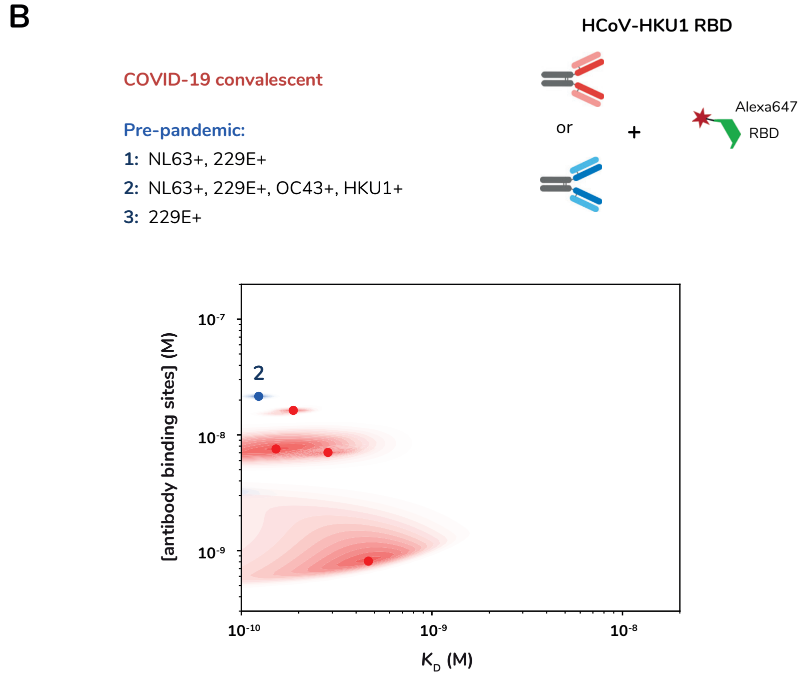


**Suppl. Figure 1.** Microfluidic antibody-affinity profiling against HCoV-HKU1 spike S1 and RBD reveal high affinity antibodies at even lower concentrations than against HCoV-NL63. **(A)** and **(B)** Probability density plots of MAAP against fluorescently labeled HCoV-HKU1 spike S1 **(A)** and HCoV-HKU1 RBD **(B)**. The graphs show the affinity ($K_D$) and the molar concentration of antibody binding sites for each of the convalescent COVID-19 sera (red) and the pre-pandemic sera (blue) where pre-pandemic serum **2** is seropositive for HCoV-HKU1. Points correspond to the maximum a posteriori values in the two-dimensional posterior probability distribution, and shaded regions to the probability density.

#### MAAP results against SARS-CoV-2 spike antigens: RBD, S1 and S2.

| SARS-CoV-2 RBD | | | | | | |
|---|---|---|---|---|---|---|
| | [Antibody binding sites] /M | | | Kd /M | | |
| **Sample** | **Best fit** | **Lower CI** | **Upper CI** | **Best fit** | **Lower CI** | **Upper CI** |
| Convalescent 1 | 3.90E-08 | 8.11E-09 | 6.58E-08 | 6.58E-09 | 3.90E-10 | 2.08E-08 |
| Convalescent 2 | 9.00E-08 | 7.30E-10 | 1.52E-06 | 1.11E-08 | 1.87E-10 | 1.87E-06 |
| Convalescent 3 | 1.00E-07 | 3.51E-10 | 2.08E-06 | 1.23E-09 | 1.11E-10 | 1.52E-06 |
| Convalescent 4 | 1.69E-07 | 1.23E-07 | 2.08E-07 | 6.58E-09 | 3.51E-09 | 1.00E-08 |
| Convalescent 5 | 2.31E-07 | 1.52E-07 | 2.85E-07 | 5.34E-09 | 2.08E-09 | 9.00E-09 |
| Convalescent 6 | 2.31E-08 | 1.52E-08 | 2.85E-08 | 1.23E-09 | N/A | 3.90E-09 |
| Convalescent 7 | 1.69E-08 | 8.11E-09 | 4.33E-08 | 2.31E-08 | 4.33E-09 | 3.90E-08 |
| Convalescent 8 | 3.51E-08 | 1.23E-08 | 5.33E-08 | 8.11E-09 | 5.34E-10 | 2.56E-08 |
| Convalescent 9 | 7.30E-08 | 9.00E-10 | 1.69E-06 | 2.85E-08 | 2.85E-10 | 2.85E-06 |
| Pre-Pandemic 1 | N/A | N/A | N/A | N/A | N/A | N/A |
| Pre-Pandemic 2 | N/A | N/A | N/A | N/A | N/A | N/A |
| Pre-pandemic 3 | N/A | N/A | N/A | N/A | N/A | N/A |

| SARS-CoV-2 S1 | | | | | | |
|---|---|---|---|---|---|---|
| | [Antibody binding sites] /M | | | Kd /M | | |
| **Sample** | **Best fit** | **Lower CI** | **Upper CI** | **Best fit** | **Lower CI** | **Upper CI** |
| Convalescent 1 | 1.87E-08 | 1.00E-09 | 3.90E-07 | N/A | N/A | 1.23E-06 |
| Convalescent 2 | N/A | N/A | N/A | N/A | N/A | N/A |
| Convalescent 3 | N/A | N/A | N/A | N/A | N/A | N/A |
| Convalescent 4 | 1.00E-07 | 8.11E-08 | 1.23E-07 | 1.37E-09 | 5.92E-10 | 2.85E-09 |
| Convalescent 5 | 4.80E-07 | 3.51E-07 | 5.33E-07 | 1.37E-08 | 8.11E-09 | 2.08E-08 |
| Convalescent 6 | 1.52E-08 | 2.31E-09 | 1.52E-08 | N/A | N/A | 3.90E-09 |
| Convalescent 7 | 1.87E-08 | 9.00E-10 | 2.85E-08 | N/A | N/A | 1.00E-08 |
| Convalescent 8 | N/A | N/A | N/A | N/A | N/A | N/A |
| Convalescent 9 | N/A | N/A | N/A | N/A | N/A | N/A |
| Pre-Pandemic 1 | N/A | N/A | N/A | N/A | N/A | N/A |
| Pre-Pandemic 2 | N/A | N/A | N/A | N/A | N/A | N/A |
| Pre-pandemic 3 | N/A | N/A | N/A | N/A | N/A | N/A |

| SARS-CoV-2 S2 | | | | | | |
|---|---|---|---|---|---|---|
| | [Antibody binding sites] /M | | | Kd /M | | |
| Sample | Best fit | Lower CI | Upper CI | Best fit | Lower CI | Upper CI |
| Convalescent 1 | 5.33E-08 | 4.33E-08 | 6.58E-08 | N/A | N/A | 8.11E-10 |
| Convalescent 2 | 1.87E-08 | 4.33E-09 | 2.31E-08 | 1.52E-09 | 1.37E-10 | 8.11E-09 |
| Convalescent 3 | 1.23E-08 | 9.00E-09 | 2.08E-08 | 7.30E-09 | 2.31E-09 | 1.11E-08 |
| Convalescent 4 | 9.00E-08 | 6.58E-08 | 1.00E-07 | 1.52E-09 | 5.34E-10 | 2.85E-09 |
| Convalescent 5 | 3.51E-07 | 2.31E-07 | 3.90E-07 | 3.90E-09 | 1.69E-09 | 5.34E-09 |
| Convalescent 6 | 3.16E-08 | 2.31E-08 | 3.90E-08 | 1.23E-09 | 1.23E-10 | 3.16E-09 |
| Convalescent 7 | 1.11E-07 | 8.11E-08 | 1.37E-07 | 3.51E-09 | 1.23E-09 | 5.34E-09 |
| Convalescent 8 | 1.52E-07 | 1.00E-07 | 1.69E-07 | 1.00E-09 | 1.69E-10 | 2.08E-09 |
| Convalescent 9 | 1.52E-08 | 6.58E-09 | 1.69E-08 | N/A | N/A | 3.51E-09 |
| Pre-Pandemic 1 | N/A | N/A | N/A | N/A | N/A | N/A |
| Pre-Pandemic 2 | N/A | N/A | N/A | N/A | N/A | N/A |
| Pre-pandemic 3 | N/A | N/A | N/A | N/A | N/A | N/A |

**Suppl. Table 1.** Best fit values and confidence intervals for all the MAAP measurements against the SARS-CoV-2 spike antigens RBD, S1 and S2. Highlighted in red: no binding could be determined. Highlighted in yellow: The data do not provide information on a lower bound on $K_D$ nor, in some cases, a peak in distribution. The upper CI given corresponds to the approximate upper bound on $K_D$.

#### MAAP results against HCoV-NL63 spike antigens: RBD and S1.

| HCoV-NL63 S1 | | | | | | |
|---|---|---|---|---|---|---|
| | [Antibody binding sites] /M | | | Kd /M | | |
| Sample | Best fit | Lower CI | Upper CI | Best fit | Lower CI | Upper CI |
| Convalescent 1 | 2.95E-08 | 2.23E-08 | 4.04E-08 | N/A | N/A | 1.42E-09 |
| Convalescent 2 | 4.04E-08 | 2.95E-08 | 5.52E-08 | N/A | N/A | 2.06E-09 |
| Convalescent 3 | 4.48E-08 | 3.05E-08 | 6.81E-08 | N/A | N/A | 2.06E-09 |
| Convalescent 4 | 2.66E-08 | 1.87E-08 | 3.63E-08 | N/A | N/A | 1.07E-09 |
| Convalescent 5 | 2.66E-08 | 1.94E-08 | 3.51E-08 | N/A | N/A | 1.35E-09 |
| Convalescent 6 | 5.52E-08 | 4.18E-08 | 7.56E-08 | N/A | N/A | 1.56E-09 |
| Convalescent 7 | 5.92E-08 | 4.64E-08 | 7.05E-08 | N/A | N/A | 4.23E-10 |
| Convalescent 8 | 3.16E-08 | 1.94E-08 | 4.04E-08 | N/A | N/A | 1.71E-09 |
| Convalescent 9 | 2.85E-08 | 2.23E-08 | 4.18E-08 | N/A | N/A | 1.87E-09 |
| Pre-Pandemic 1 | 3.27E-08 | 2.39E-08 | 4.98E-08 | N/A | N/A | 1.87E-09 |
| Pre-Pandemic 2 | 9.00E-08 | 6.13E-08 | 1.28E-07 | 1.29E-09 | 1.38E-10 | 3.27E-09 |
| Pre-pandemic 3 | 5.52E-08 | 2.66E-08 | 8.40E-08 | 3.59E-09 | 2.92E-10 | 7.92E-09 |

| HCoV-NL63 RBD | | | | | | |
|---|---|---|---|---|---|---|
| | [Antibody binding sites] /M | | | Kd /M | | |
| Sample | Best fit | Lower CI | Upper CI | Best fit | Lower CI | Upper CI |
| Convalescent 1 | 2.95E-08 | 2.39E-08 | 3.51E-08 | 1.23E-09 | 2.66E-10 | 3.27E-09 |
| Convalescent 2 | 2.85E-08 | 2.48E-08 | 3.39E-08 | N/A | N/A | 4.43E-10 |
| Convalescent 3 | 3.39E-08 | 2.15E-08 | 4.33E-08 | 2.98E-09 | 5.34E-10 | 7.92E-09 |
| Convalescent 4 | 2.56E-08 | 2.15E-08 | 2.85E-08 | N/A | N/A | 3.05E-10 |
| Convalescent 5 | 3.05E-08 | 2.48E-08 | 3.90E-08 | N/A | N/A | 2.06E-09 |
| Convalescent 6 | 3.90E-08 | 3.39E-08 | 4.33E-08 | N/A | N/A | 3.68E-10 |
| Convalescent 7 | 3.27E-08 | 2.85E-08 | 3.63E-08 | N/A | N/A | 5.09E-10 |
| Convalescent 8 | 2.66E-08 | 1.75E-08 | 3.39E-08 | 2.60E-09 | 1.23E-09 | 3.94E-09 |
| Convalescent 9 | 1.75E-08 | 1.52E-08 | 2.39E-08 | N/A | N/A | 1.79E-09 |
| Pre-Pandemic 1 | 2.85E-08 | 2.48E-08 | 3.27E-08 | 4.43E-10 | 1.75E-10 | 8.11E-10 |
| Pre-Pandemic 2 | 3.51E-08 | 2.95E-08 | 3.90E-08 | 5.09E-10 | 1.15E-10 | 1.23E-09 |
| Pre-pandemic 3 | 2.23E-08 | 2.01E-08 | 2.48E-08 | N/A | N/A | 1.92E-10 |

**Suppl. Table 2.** Best fit values and confidence intervals for all the MAAP measurements against the HCoV-NL63 spike antigens RBD and S1. Highlighted in yellow: The data do not provide information on a lower bound on $K_D$ nor, in some cases, a peak in distribution. The upper CI given corresponds to the approximate upper bound on $K_D$.

#### MAAP results against HCoV-HKU1 spike antigens: RBD and S1.

| HCoV-HKU1 S1 | | | | | | |
|---|---|---|---|---|---|---|
| | [Antibody binding sites] /M | | | Kd /M | | |
| **Sample** | **Best fit** | **Lower CI** | **Upper CI** | **Best fit** | **Lower CI** | **Upper CI** |
| Convalescent 3 | 2.08E-08 | 1.18E-08 | 2.65E-08 | 3.12E-09 | 1.63E-09 | 5.49E-09 |
| Convalescent 4 | 2.75E-09 | 3.65E-10 | 3.81E-09 | 3.24E-10 | 1.08E-11 | 3.97E-09 |
| Convalescent 5 | 9.27E-09 | 6.18E-09 | 1.28E-08 | 4.13E-10 | 7.55E-11 | 1.18E-09 |
| Convalescent 6 | 1.09E-08 | 6.70E-09 | 2.87E-08 | 7.00E-09 | 2.65E-09 | 1.23E-08 |
| Convalescent 7 | 3.66E-08 | 2.45E-08 | 5.06E-08 | 1.63E-09 | 6.70E-10 | 2.87E-09 |
| Convalescent 8 | 4.85E-09 | 1.70E-09 | 1.63E-08 | 5.06E-09 | 6.70E-10 | 1.23E-08 |
| Convalescent 9 | 7.05E-09 | 5.72E-09 | 8.11E-09 | N/A | N/A | 3.51E-11 |
| Pre-Pandemic 1 | 1.28E-08 | 6.70E-09 | 1.77E-08 | 1.51E-09 | 6.70E-10 | 2.87E-09 |
| Pre-Pandemic 2 | 9.67E-08 | 6.45E-08 | 1.23E-07 | 4.67E-09 | 2.65E-09 | 6.45E-09 |
| Pre-pandemic 3 | N/A | N/A | N/A | N/A | N/A | N/A |

| HCoV-HKU1 RBD | | | | | | |
|---|---|---|---|---|---|---|
| | [Antibody binding sites] /M | | | Kd /M | | |
| **Sample** | **Best fit** | **Lower CI** | **Upper CI** | **Best fit** | **Lower CI** | **Upper CI** |
| Convalescent 4 | 3.05E-09 | 4.98E-10 | 3.51E-09 | 2.85E-11 | 1.07E-11 | 9.33E-10 |
| Convalescent 6 | 7.05E-09 | 5.72E-09 | 8.11E-09 | 2.85E-10 | 1.42E-10 | 4.64E-10 |
| Convalescent 7 | 1.63E-08 | 1.42E-08 | 1.75E-08 | 1.75E-10 | 1.15E-10 | 2.48E-10 |
| Convalescent 8 | 7.05E-09 | 4.98E-09 | 1.15E-08 | 1.23E-10 | 1.07E-11 | 5.72E-10 |
| Convalescent 9 | 7.05E-09 | 5.72E-09 | 8.11E-09 | N/A | N/A | 3.51E-11 |
| Pre-Pandemic 1 | 3.76E-09 | 3.27E-09 | 3.76E-09 | N/A | N/A | 5.34E-11 |
| Pre-Pandemic 2 | 2.15E-08 | 2.01E-08 | 2.15E-08 | | 8.70E-11 | 1.52E-10 |
| Pre-pandemic 3 | 3.27E-09 | 2.66E-09 | 3.76E-09 | N/A | N/A | 1.32E-10 |

**Suppl. Table 3.** Best fit values and confidence intervals for all the MAAP measurements against the HCoV-HKU1 spike antigens RBD and S1. Highlighted in red: no binding could be determined. Highlighted in yellow: The data do not provide information on a lower bound on $K_D$ nor, in some cases, a peak in distribution. The upper CI given corresponds to the approximate upper bound on $K_D$.
