## Supplementary Material_QC for "Understanding the role of memory re-activation and cross-reactivity in the defense against SARS-CoV-2"

**Suppl. Material 1.** Quality Control (QC) pipeline.

### <u>Quality control (QC) pipeline and dataset overview</u>

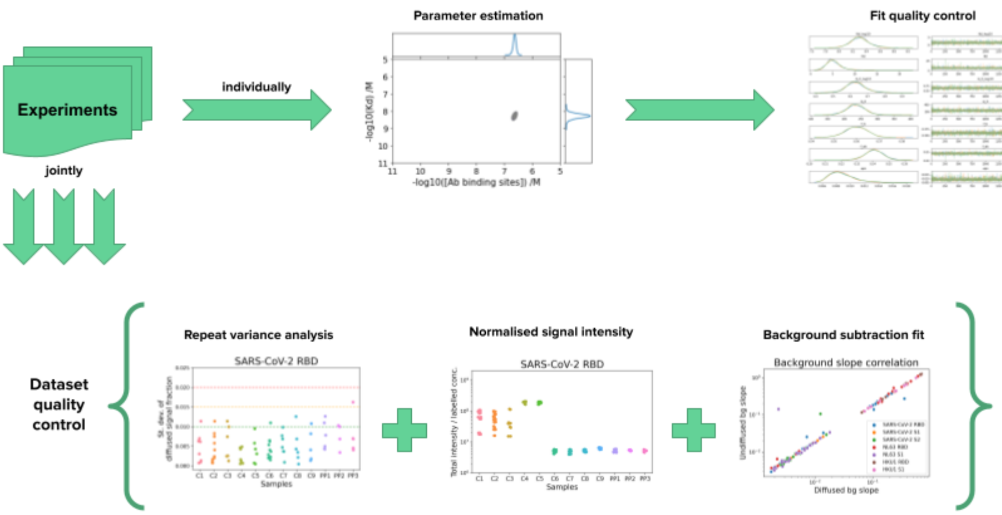


**Figure 1.** An overview of the QC pipeline. Individual samples are processed to fit model parameters. The quality of these fits can be assessed by analyzing diagnostic trace plots for the Markov Chain Monte Carlo sampling procedure (top row). By combining related samples, further QC tests can identify outlier points or problems within the dataset (bottom row).

| | No serum | | Individual 1 | | Individual 2 | |
|---|---|---|---|---|---|---|
| **No labeled antigen** | | | Serum 1 background | | Serum 2 background | |
| | | | Day 1 | Day 2 | Day 1 | Day 2 |
| **Antigen 1** | Free labeled antigen 1 | Week 1 | Experiment 1-1 part 1 | Experiment 1-1 part 2 | | |
| | | Week 2 | | | Experiment 2-1 part 1 | Experiment 2-1 part 2 |
| **Antigen 2** | Free labeled antigen 2 | Week 1 | Experiment 1-2 part 1 | Experiment 1-2 part 2 | | |
| | | Week 2 | | | Experiment 2-2 part 1 | Experiment 2-2 part 2 |

**Table 1.** An overview of how measurements in the dataset relate to each other. A simplified example is shown, where two labeled antigens and sera from two individuals are considered. “Experiment 1-2” refers to serum sample 1 measured against antigen 2. See text for a detailed explanation of the structure of the dataset and relationships between measurements.

As part of constructing the dataset used in this research, we developed a series of procedures that we jointly refer to as "quality control (QC) pipeline". Their purpose is manifold. Firstly, they serve as simple sanity checks to ensure that data processing and model fitting was performed correctly. Secondly, they help identify problematic measurements, samples, or model fits. Based on this, decisions to re-collect or exclude parts of the data can be made. Finally, if assumptions about relationships in the dataset are appropriately tested, information can be shared between experiments and samples in a robust way, which improves model fitting.

It is important to note that we do not include instrument-intrinsic errors, nor operator errors, in this discussion. We assume that the data collected is reliable to the extent that all input species, samples, and concentrations are correctly labeled, and no measurements flagged by the instrument as errors have been included in the dataset.

The structure of the dataset is presented in **Table 1**. Sera from 12 individuals were measured against five labeled antigens. A "data point" or "measurement" refers to a specific combination of labeled antigen concentration and serum dilution. These were typically performed in replicates (triplicates). A set of measurements relating to one serum against one antigen, with associated controls, is referred to as an "experiment" (i.e., there were 12x5=60 experiments). Model fitting and parameter estimation was performed at the level of an individual experiment. The controls consisted of measurements of labeled antigens in PBS buffer (no serum) and of serum background fluorescence in the absence of labeled antigens at varying serum dilutions (see "Materials and Methods", Microfluidic antibody-affinity profiling of serum antibodies (MAAP)). No-serum controls were typically performed once per week and used for all experiments with that specific labeled antigen during the week. Each serum was analyzed across 1–4 days. Serum background fluorescence measurements were typically performed each day and were used for background correction of data points measured on the same day. Therefore, for each antigen there will be several no-serum measurement sets, and each serum sample will have multiple background measurement sets.

**Figure 1** summarizes the components of the QC pipeline, each addressed in their own section below in detail. Firstly, we analyze the model fit quality of each experiment through diagnostic trace plots for the Markov Chain Monte Carlo sampling procedure. Then, we pull experiments together into a combined dataset and look at repeat variance, background subtraction fits and normalized fluorescence intensity.

### Model fit QC

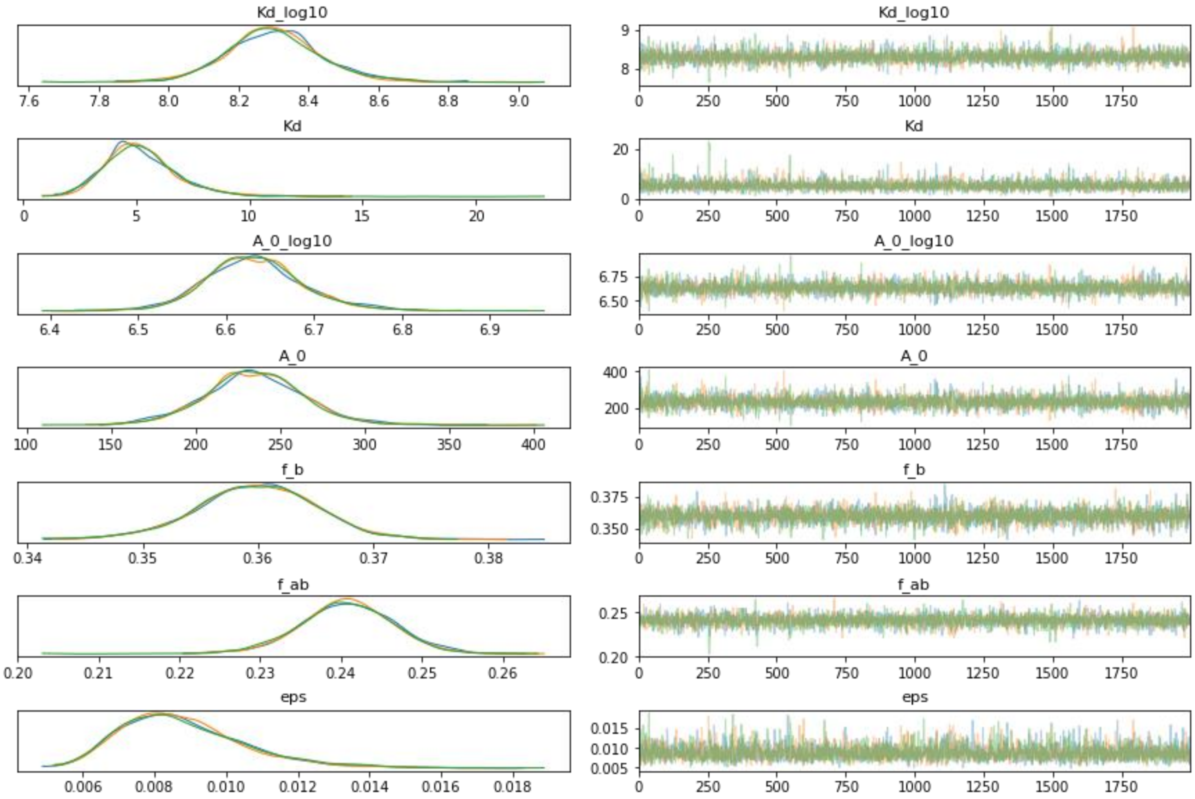


**Figure 2.** An example trace plot generated by ArviZ (https://arviz-devs.github.io/arviz/) "plot trace" function. The left column shows kernel density estimates of the marginal posterior distributions of all model parameters fitted using Markov Chain Monte Carlo. The right column shows individual sampled parameter values at each step during the sampling procedure. Individual chains are shown in different colors in both columns. Black rug plot indicates divergences.

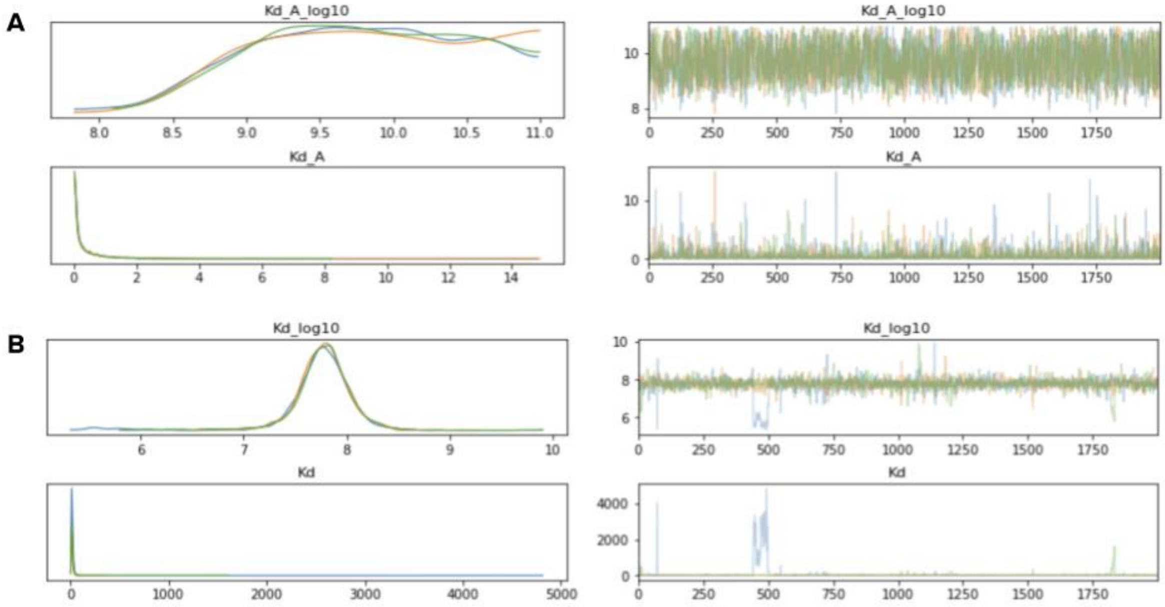


**Figure 3.** Examples of pathologies arising during model fitting. Unconverged parameter estimates **(A)**. Meta-stable divergences **(B)**.

Model parameter fitting is performed in a Bayesian statistical framework with Markov Chain Monte Carlo sampling using Python packages PyMC3 (https://docs.pymc.io/) and ArviZ (https://arviz-devs.github.io/arviz/). We rely on the standard diagnostic procedure to assess model fit quality and identify low-quality fits - trace plot analysis, an example of which is shown in **Figure 2**.

The left column of this plot shows kernel density estimates of the marginal posterior distributions for all model parameters. A good model fit will have stationary and well-converged marginal posteriors ("tight" approximately bell-shaped curves). Furthermore, a good model fit will display concordance between different sampled chains (shown in different colors). The right column shows individual sampled parameter values at each step during the sampling procedure. A good model fit will display low variance and no (or very few) divergences.

Pathologies that may arise include unconverged parameter estimates (**Figure 3A**, showing a case where the marginal posterior distribution is bounded only on one side but not the other); or meta-stable divergences (**Figure 3B**), i.e. where a chain switches from oscillating around one mean value of a parameter to a different mean value for a contiguous block of sampled values.

It is important to visually assess the quality of each model fit in order to decide whether the parameter estimates can be trusted.

### Repeat variance analysis 1

The design of control measurements for a complicated dataset needs to take into account various potential sources of variability. However, not all of these will indeed result in significant variation. For example, it is prudent to test the time-dependent change in labeled antigen signal without serum. Re-measuring it weekly allows for a new set of no-serum control measurements to be used with samples measured during that week. However, this comes at a cost of increasing the total number of measurements that need to be performed and reducing the effective number of data points used in the analysis of each sample.

As was shown in **Table 1**, free labeled antigen measurements (without serum) were performed in weekly batches. These measurement points contribute the most to fixing the free labeled species size in the model fitting step. If the assumption holds that there is no week-to-week variation in the free labeled species measurements, we could pull all those replicates together and obtain a more reliable estimate of their size. (While we work with diffused signal fraction in our analysis, it is directly related to species size.)

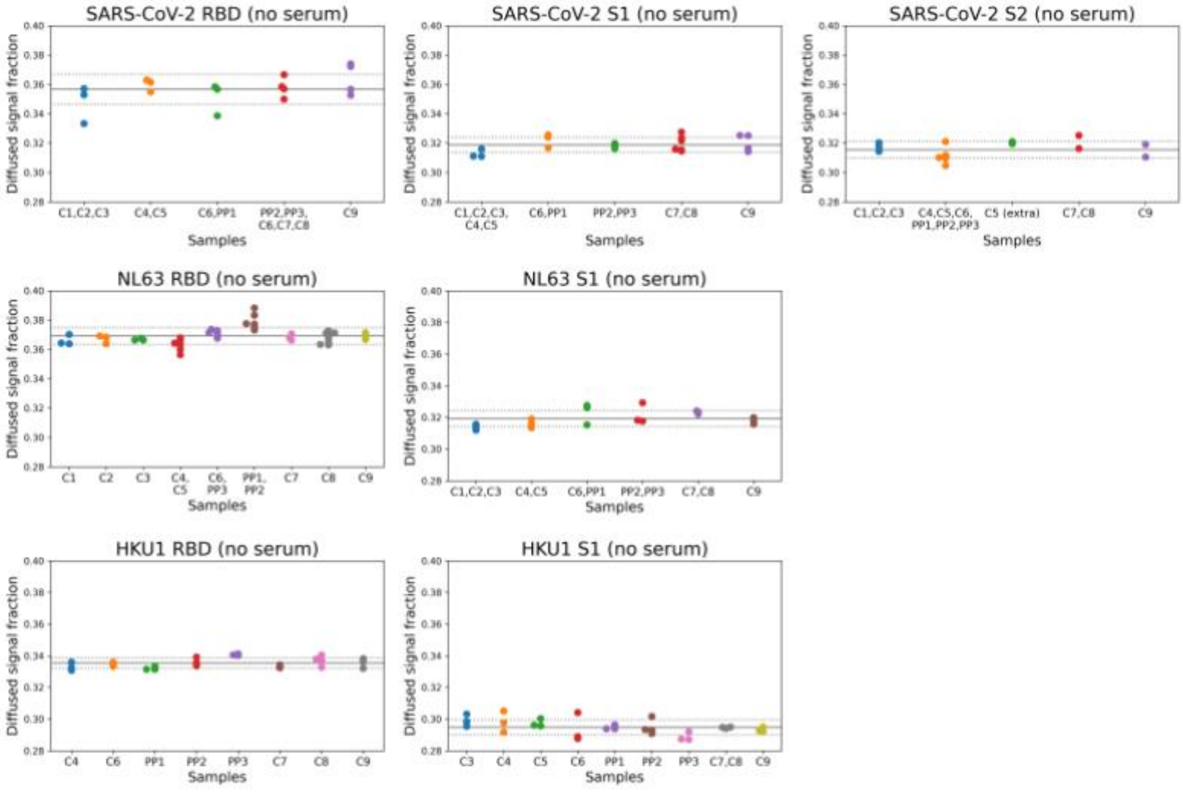


**Figure 4.** Repeat variance analysis of free labeled antigen measurements (without serum). Diffused signal fraction corresponds to the fraction of fluorescence intensity signal detected in the diffused half of the channel, out of total fluorescence intensity detected. Solid line shows the mean value, with dotted lines showing 1 standard deviation above and below the mean value. Convalescent sera (**C1–9**) and pre-pandemic sera (**PP1–3**) are grouped on the x-axis according to the weekly batches in which they were measured together. For each batch, a separate no-serum measurement set was performed using the free labeled antigen. The week-to-week variation between these is not significant.

**Figure 4** compares the weekly free labeled species measurements for all antigens. For all antigens in this analysis, there was no significant week-to-week variation or time-dependent change in signal. Therefore, instead of using only the weekly batch in each sample's model fit, we pooled all measurements for a single antigen together, in order to provide a more precise estimate of the free labeled species size.

### Repeat variance analysis 2

The purpose of this dataset-level QC step is to detect outlier measurements that may need to be removed in order to improve the quality of the model fitting. We observe that performing this analysis on the level of the whole dataset allows for a more robust comparison between replicates, as opposed to evaluating individual experiments on their own.

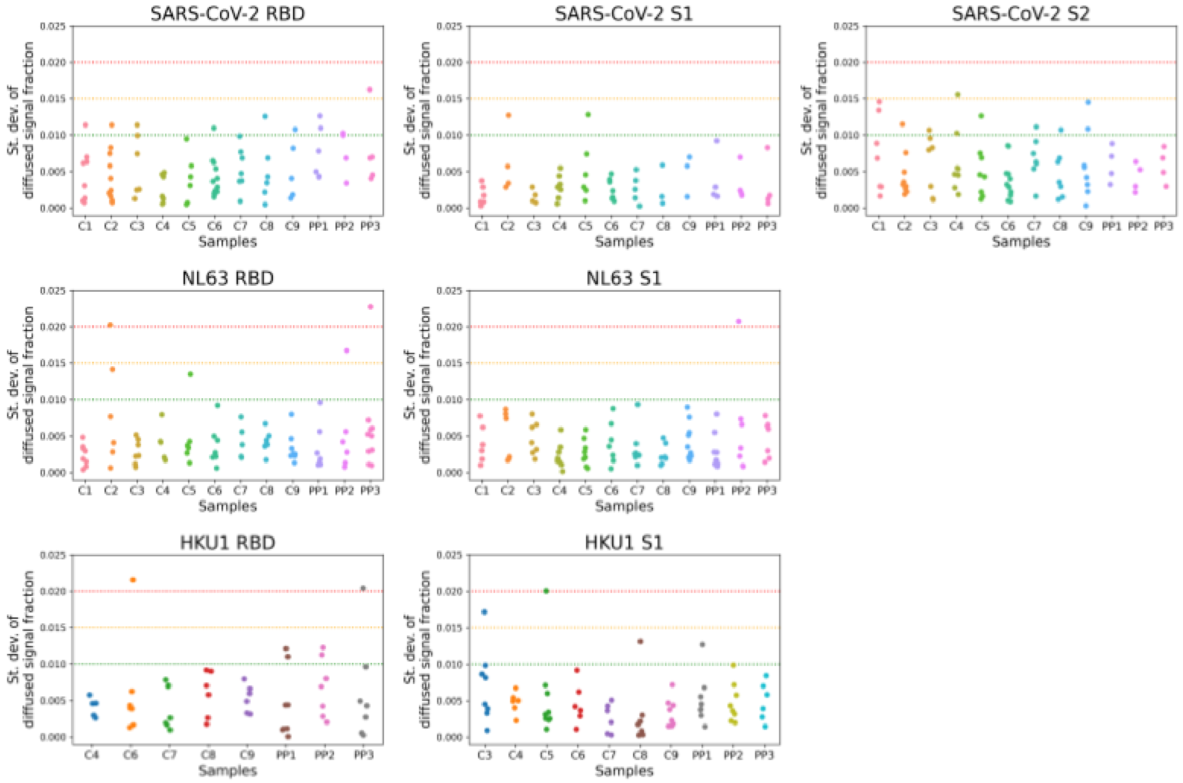


**Figure 5**. Standard deviations of replicate measurements. Diffused signal fraction corresponds to the fraction of fluorescence intensity signal detected in the diffused half of the channel, out of total fluorescence intensity detected. For each measurement performed in replicate, the standard deviation is computed. The horizontal lines represent empirical cut-off values. Measurements falling below the green line (0.010) are ideal, between the green and the yellow line (0.015) are acceptable, between the yellow and the red line (0.020) are worrying, and above the red line are almost certainly problematic. Only a small number of measurements fall above the yellow line; these require manual curation and potential exclusion from the dataset as outliers.

As all measurements are performed in replicates, one way to detect outliers is to consider the standard deviation of signal within a replicate set. This is plotted for all samples in the dataset in **Figure 5**. From other datasets obtained on the same platform (data not shown), we have established empirical guidelines for acceptable levels of variance. These are visualized as colored horizontal lines in the figure. Variance above acceptable levels can lead to unstable model fits or even lack of convergence. This analysis allows the identification and exclusion of such problematic data points. Alternatively, further measurements using the identified input concentrations can be taken until variance is reduced to acceptable levels and the model fits are improved appropriately.

As can be seen in the case of the dataset used in this work, only a very small number of replicate sets have standard deviations exceeding the yellow threshold (0.015), which we consider to be the acceptability cut-off. These replicate sets were manually reviewed and outliers with disproportionate impact on model fit quality were removed.

### Normalized signal intensity

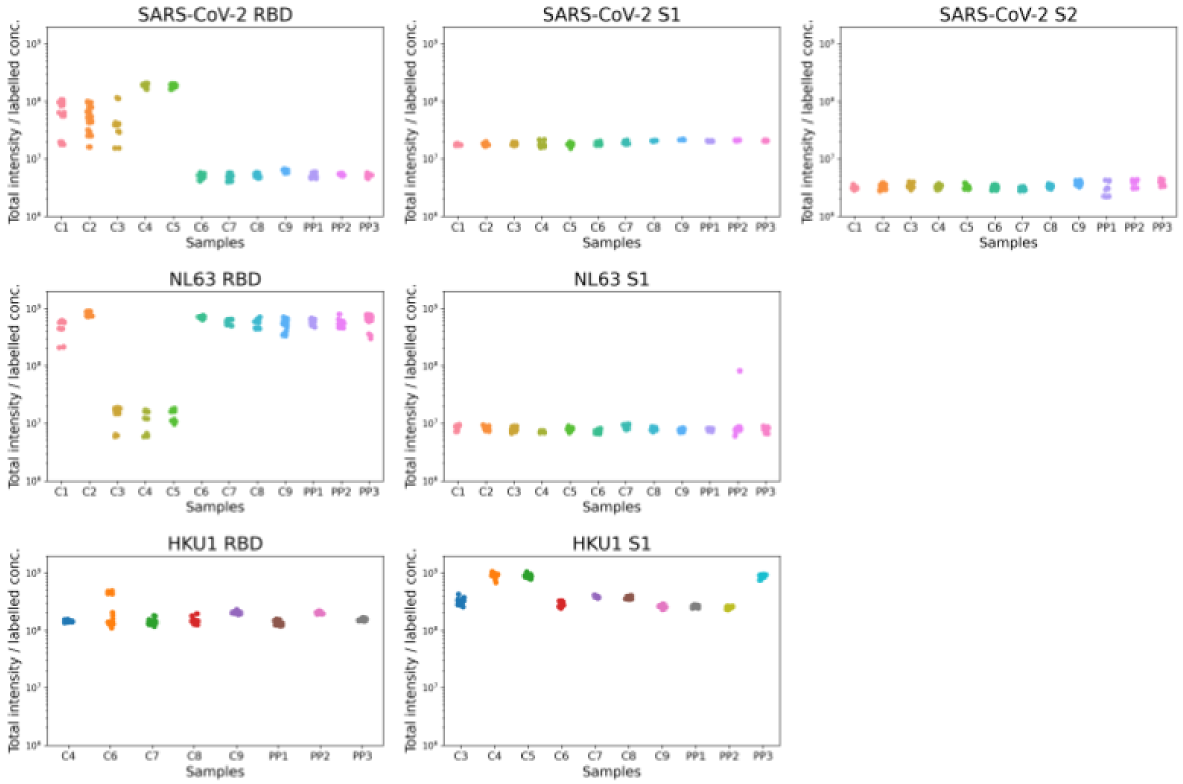


**Figure 6**. Total fluorescence intensity of each measurement, normalized by the concentration of the labelled species. Assuming a relatively uniform amount of fluorescent label per unit protein, for each protein type all measurements should tend to the same value. This is the case for all samples except SARS-CoV-2 RBD **C1–3**, **C4–5** and HCoV-NL63 RBD **C3–5**, where a different instrument was used to perform the measurements. Additionally, significant deviations within a sample, such as the outlier measurement in HCoV-NL63 S1 **PP2** and the spread of measurements in SARS-CoV-2 RBD **C1–3** are identified as causes for concern that need to be addressed.

An essential assumption that is being made as part of the data analysis is that the labeled species are uniformly labeled with a fluorescent tag, as the observed signal intensity is taken to be a direct measurement of the concentration of said species. Therefore, validating this assumption is an important part of quality control.

In the simplest case, with no effect from confounding factors, if the total measured intensity (across both the diffused and the undiffused halves of the channel) is divided by the concentration of the labeled species, for each protein type all measurements should tend to the same value, which is the amount of fluorescent label per unit protein. (Note that this will be different between different proteins, as their labeling efficiencies will differ.) As shown in **Figure 6**, for most samples, the assumption holds with most measurements for a protein type lining up.

However, care needs to be taken to account for confounding factors that may affect the interpretation of this analysis. For example, experiments SARS-CoV-2 RBD **C1–3**, **C4–5** and HCoV-NL63 RBD **C3–5** were measured on different instruments from the rest of the SARS-CoV-2 and HCoV-NL63 experiments. This analysis revealed that the total fluorescence intensities cannot be directly compared between samples taken on different instruments. (See also *Background subtraction fit* section below.)

However, within each group, the normalized intensity value distributions were similar. Thus, measurements from different instruments cannot be pooled for analyzing one sample but as long as separate samples are measured on separate instruments, the difference in total fluorescence intensity is not an issue.

This analysis reveals measurements that are concerning. For example, an outlier is identified in the HCoV-NL63 S1 **PP2** sample. Interestingly, this outlier is also the data point responsible for the abnormal variance of a replicate set identified in *Repeat variance analysis 2* section (see **Figure 5**). Furthermore, the wide distribution of normalized intensity values in samples such as SARS-CoV-2 RBD **C1–3** are indicative of a potential problem with those samples and require further investigation. It was possible to establish that in most cases where this occurred, the likely source of variation was due to using different preparations of labeled protein. While this variation cancels out in the model fitting step (the ratio of intensity values in the two halves of the channel remains constant, and it is the ratio which is used), care needs to be taken to ensure that such sources of variation are recognized and correctly accounted for if these data are to be used in other ways, e.g. for sharing information between experiments. In this case, only measurements performed using the same labeled protein preparation must be used.

More generally, analyses like these highlight the importance of rigorously recording metadata associated with each measurement. Information about sample preparation, instrument and user IDs, and even timestamps of experiments are just some of the potential sources of systematic variation, the effects of which - while cancelling out in some instances - can lead to potential errors or wrong conclusions being drawn from the data. In the next section, we describe one more example of such “cryptic” source of variation.

### Background subtraction fit

When working with serum samples, it is crucial to account for intrinsic fluorescence and subtract that from the measured intensities (see "Materials and Methods", Microfluidic antibody-affinity profiling of serum antibodies (MAAP)). In the current dataset, background estimation for each serum sample was performed multiple times, as described in **Table 1**. This provides the opportunity to not only compare the background fits between different sera samples but also to check for any variation within each serum. We perform this analysis by focusing on the slope of the background fluorescence fit (intensity versus serum dilution, forced through the origin) and on the residuals of that linear fit.

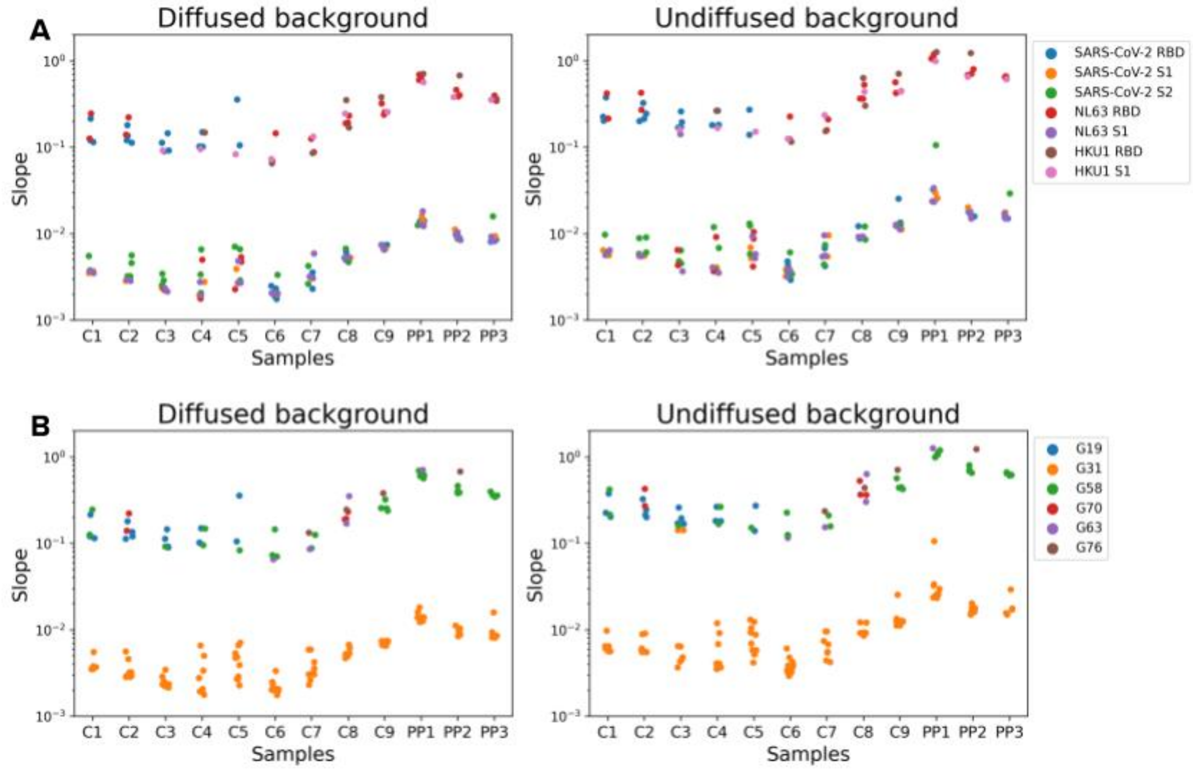


**Figure 7.** The slopes of linear fits for background serum fluorescence estimation, per sample, for both diffused and undiffused halves of the channel. The same data points are color-coded by labeled species **(A)** or instrument ID **(B)**. The expectation is for all slopes within a serum sample to tend to the same value, if there are no sources of variation between individual background fits. This assumption, however, is violated, with measurements on the G31 instrument consistently resulting in lower slope estimates.

Firstly, we tested the assumption that the background fits performed on the same serum sample multiple times should all tend to the same value. This value may be different between different sera, as their individual composition will result in different levels of background fluorescence, but the same serum measured multiple times should produce the same background fit. As **Figure 7** demonstrates, this is not the case for any of the 12 sera, neither in the diffused nor in the undiffused half of the channel.

Having observed this effect, we set out to find the systematic difference between the individual measurements which would account for this variation. Splitting the data by the type of labeled protein for which the background fits were performed (**Figure 7A**) is a proxy for time-dependent variation and batch effects, as experiments were typically performed with one labeled protein at a time. However, this did not explain the variation. Instead, splitting the data by the instrument ID on which the measurements were performed (**Figure 7B**) fully accounted for the differences in slope. It appears that the instrument G31 consistently measured lower absolute intensity values than other instruments used in this work. (This also accounts for the differences in normalized signal intensity observed in **Figure 5**.)

In this case, the lower absolute intensity values are not affecting the interpretation of the data as long as the effects between the diffused and the undiffused halves of the channel are consistent with each other and cancel out. As discussed earlier (in the *Normalized signal intensity* section), this variation may, however, be important in other cases. Therefore, this analysis further underscores the importance of recording and analyzing the variation arising due to systematic differences between individual measurements, such as the instrument ID.

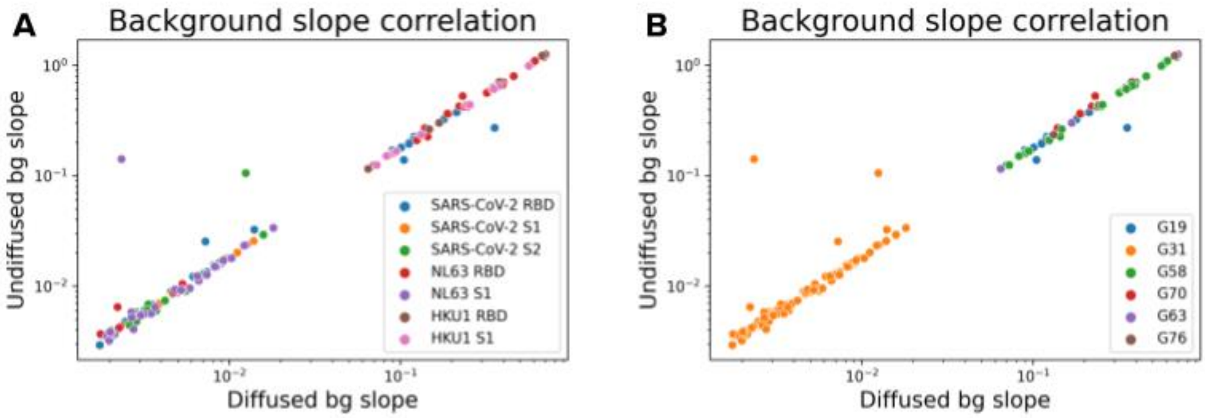


**Figure 8.** Correlation of the background fit slopes between the diffused and the undiffused halves of the channel. The same data points are color-coded by labeled species **(A)** or instrument ID **(B)**. Ideally, all points should fall on the y=x line, as that would imply consistency between the intrinsic fluorescence of the serum in both halves of the channel. Those points where that is not the case indicate potentially problematic sets of data from which the corresponding background fits were estimated - e.g. there may be significant outliers affecting the fit quality.

Secondly, and following up on the statement that we expect consistency in intrinsic serum fluorescence between the diffused and the undiffused halves of the channel, the matched pairs of undiffused and diffused background slopes were plotted against each other in **Figure 8**. Any points that fall significantly off the y=x line indicate potentially problematic sets of data from which the corresponding background fits were estimated. There were only a small number of such points and closer analysis revealed outlier measurements within those background measurement sets. These were removed before subsequent analysis and model fitting.

It is also informative to observe that despite the difference in absolute intensity values noted earlier for instrument G31, most of the background slopes estimated on it fall nicely on the y=x line, alleviating any concerns about using the data measured on this instrument in further analysis.

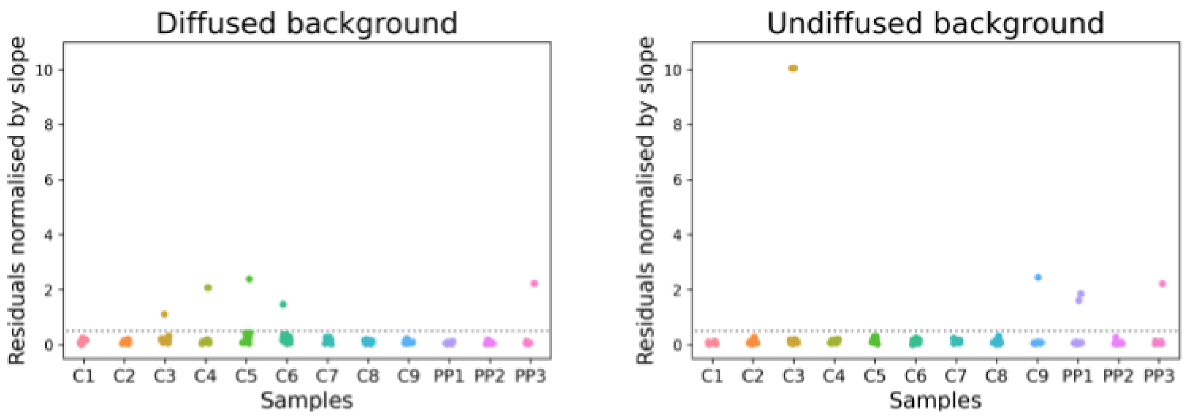


**Figure 9.** The residuals of linear fits for background serum fluorescence estimation, per sample, for both diffused and undiffused halves of the channel. The linear fit returns squared residuals, the square root of which is taken, followed by normalisation by the slope value. Points falling above the grey dotted line at 0.5 are considered to be potentially problematic - e.g. there may be significant outliers affecting the fit quality.

Finally, the residuals of the linear background fits need to be considered in order to assess the quality of the fits. For comparative purposes, the square root of the (squared) residual was normalized by the value of the corresponding slope, as shown in **Figure 9**. Only a small number of points fell above an empirically determined cut-off of 0.5. These were further investigated and were revealed to be due to outlier measurements, many of which were the same outliers as identified earlier in **Figure 8**. These were removed before subsequent analysis and model fitting.

### Conclusion

Background subtraction fit analysis concludes the series of QC tests designed for the present dataset. These are not meant to be exclusive, nor prescriptive. It is acknowledged that different datasets may contain different kinds of relationships between the individual measurements, samples, experiments and normalization or controls performed. It is important to design any QC procedure such that it tests assumptions about these relationships and tries to uncover sources of variation and potential errors or outliers in the dataset. Not only does this lead to higher quality data and more confident analysis and conclusions from such data, but it also helps design better experiments in the future.
